# Proteomic profiling of peripheral blood mononuclear cells isolated from patients with tuberculosis and diabetes copathogenesis -A pilot study

**DOI:** 10.1101/2020.05.05.078519

**Authors:** Jyoti Kundu, Shikha Bakshi, Himanshu Joshi, Sanjay K Bhadada, Indu Verma, Sadhna Sharma

## Abstract

**Background:** Diabetes is an important risk factor for developing tuberculosis. This association leads to exacerbation of tuberculosis symptoms and delayed treatment of both the diseases. Molecular mechanism and biomarkers/drug targets related to copathogenesis of tuberculosis and diabetes, however, still remains to be poorly understood. In this study, proteomics based 2D-MALDI/MS approach was employed to identify host signature proteins which are altered during copathogenesis of tuberculosis and diabetes.

**Methods:** Comparative proteome of human peripheral blood mononuclear cells (PBMCs) from healthy controls, tuberculosis and diabetes patients in comparison to comorbid diabetes and tuberculosis patients was analyzed. Gel based proteomics approach followed by in gel trypsin digestion and peptide identification by mass spectrometry was used for signature protein identification.

**Results:** Total of 18 protein spots with differential expression in TBDM patients in comparison to other groups were identified. These include Vimentin, tubulin beta chain protein, superoxide dismutase, Actin related protein 2/3 complex subunit 2, PDZ LIM domain protein, Rho-GDP dissociation inhibitor, Ras related protein Rab, dCTPpyrophosphatase 1, Transcription initiation factor TFIID subunit 12, coffilin 1, three isoforms of Peptidylprolylcis-trans isomerase A, three isoforms of Protein S100A9, Protein S100A8 and SH3 domain containing protein. These proteins belonged to four functional categories i.e. structural, cell cycle/growth regulation, signaling and intermediary metabolism.

**Conclusion:** Proteins identified to be differentially expressed in TBDM patient can act as potent biomarkers and as predictors for copathogenesis of tuberculosis and diabetes.

## Introduction

Tuberculosis continues to be the global epidemic. In 2019, about 1.5 million deaths and 10million fresh TB cases has been reported worldwide (1). Despite extensive research on the biology of *M. tuberculosis*, exact mechanism of infection and immune evasion still remains elusive. Copathogenesis with HIV and diabetes further complicates the tuberculosis control measures. Although HIV infection is the topmost risk factor for development of active tuberculosis but population attributable risk of diabetes is more than that of HIV infection (2, 3) as diabetes is known to triple the risk of developing active tuberculosis (4). As diabetes is associated with various immunological dysfunctions, diabetic patients fall prey to other co-infections like tuberculosis, melioidosis and other conventional hyperglycemia related complications like various cardiovascular disorders, retinopathy, nephropathy and many more. Diabetes mellitus affected 425 million individuals worldwide in 2017 and is predicted to reach 629 million by 2045, the time at which 80% of diabetics will be residents of economically challenged countries where active tuberculosis (TB) prevails (5, 6). Countries with highest burden of diabetes are also in the list of WHO’s tuberculosis high burden countries and India and China for example, have first two positions for the copathogenesis of TB and diabetes (7, 8). The epidemiological data for association between TB and DM is increasing significantly (9, 10), however, the current literature and evidence about the connection between TB and DM is ambiguous. Most importantly, the molecular mechanism underlying this association is difficult to interpret and pathophysiology behind the co-occurrence of TB and DM is still poorly understood.

Peripheral blood mononuclear cells (PBMCs) are the primary macrophages which later on differentiate to specific tissue macrophages that play crucial role in containment of *M. tuberculosis*. In order to understand the mechanism of association between TB and DM and to findout proteins involved in combined pathogenesis, a comprehensive analysis of whole-cell protein expression in PBMCs isolated from TB and DM patients and healthy donors was performed using gel based proteomics approach.

## Methods

### Chemicals and consumables

All reagents used were of molecular grade, protease inhibitor cocktail, Phenylmethylsulfonyl fluoride (PMSF), Proteomics grade trypsin from porcine pancreas, Iodoacetic acid (IAA), Acrylamide, bis acrylamide, ficollhistopaque density gradient and Ammonium bicarbonate were procured from Sigma-Aldrich, St. Louis, MO, USA. Sodium Dodecyl Sulfate (SDS), IPG strips, Urea, thiourea, Mineral oil, Dithiothreitol (DTT), CHAPS (3-((3-cholamidopropyl) dimethylammonio)-1-propanesulfonate) were purchased from Bio-Rad Laboratories, Inc. California, USA. For protein quantification, Pierce™ 660nm Protein Assay kit from Thermo Fisher Scientific Inc. USA was used. Mouse, Anti-AGE monoclonal antibodies for western blotting were obtained from MP Biomedicals, USA. The polyvinylidenedifluoride (PVDF) membrane (0.45μm) and dialysis tubing were purchased from Millipore Corporation, MA, USA. C-18 ZipTip columns for concentrating and desalting the peptides were purchased from Millipore, Billerica, MA, USA. Pre-stained protein ladder was procured from Real Biotech Corporation, Taipei, Taiwan. All other reagents used were of high analytical grade.

### Ethical clearance

The protocol of the study was approved by Institutional ethical committee (IEC) with ref No. INT/IEC/2016/2580.

### Patients and controls

Patients of either sex were included in the study. Written informed consent was obtained from all patients. Population for study was divided into four groups (Table 1). First group consisted of healthy individuals, second group was having naïve tuberculosis and no diabetes, third group included patients having diabetes alone and fourth was confirmed cases of concurrent tuberculosis and diabetes patients diagnosed and labeled as tubercular as well as diabetic by the concerned physician attending medical and chest clinic of new OPD, Post Graduate Institute Medical Education Research, Chandigarh, India. Diagnosis was done by patient history, Montoux test, chest X-ray, sputum for AFB, or microscopic examination for tuberculosis. Patients were considered to have diabetes if they were taking an oral hypoglycemic agent or receiving insulin or were found to have two or more fasting blood glucose levels greater than 126mg/dl.

**Table 1:**
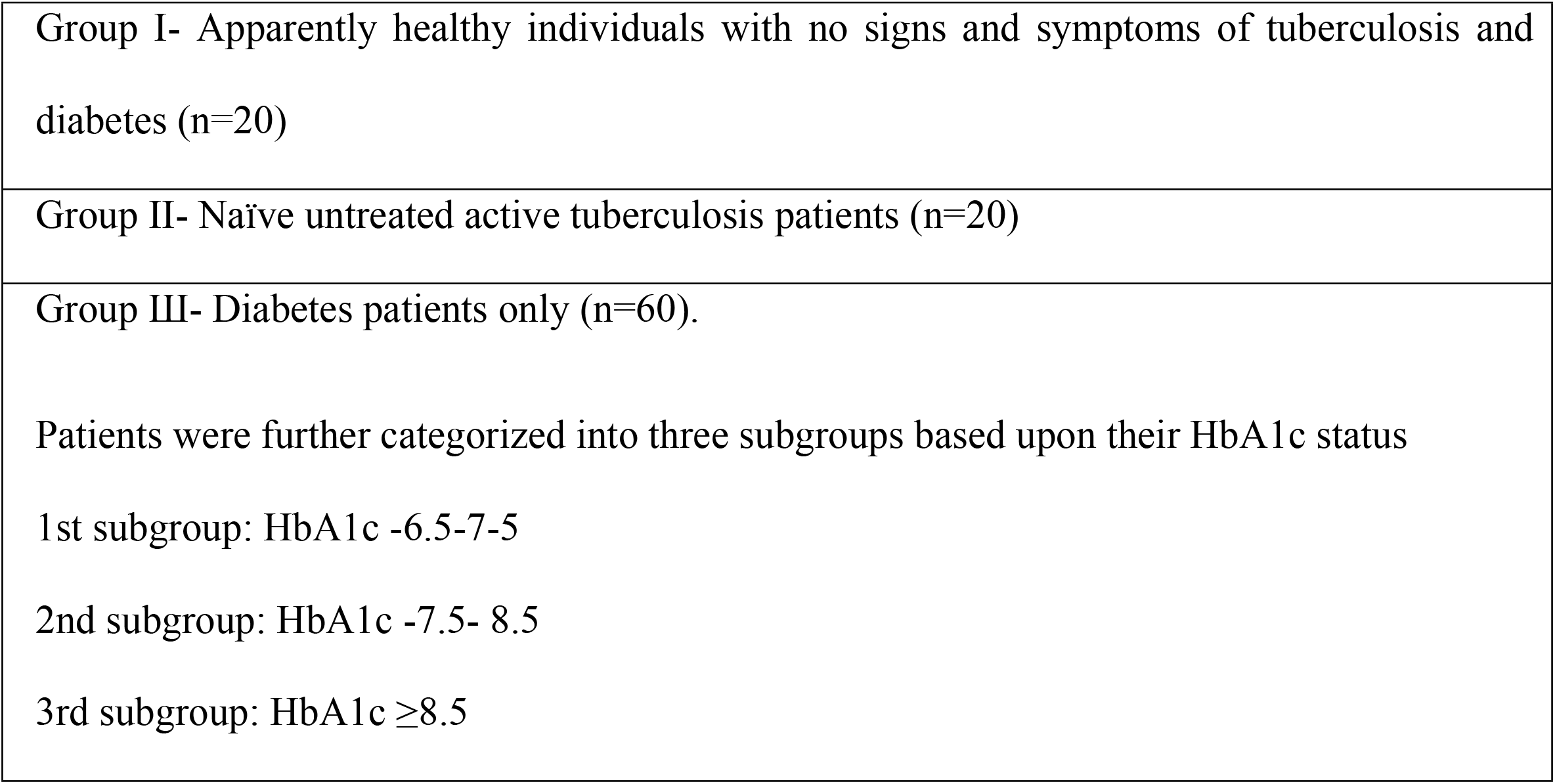
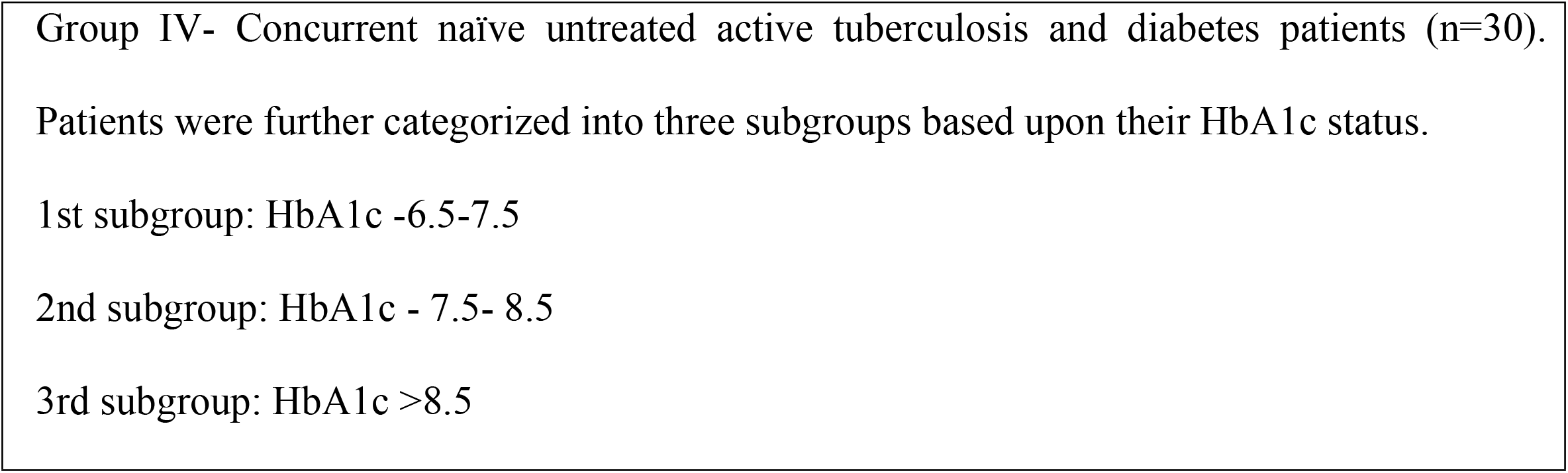
Study groups.

### Exclusion criteria

Patients with cardiovascular disease, arthritis, viral hepatitis, cancer, human immunodeficiency virus (HIV) infection, pregnancy, tuberculosis (TB) patients on anti-tubercular therapy and failure to give written informed consent were excluded from the study.

### PBMCs isolation and characterization

After confirmation of the disease status and taking the informed consent, 10mL of venous blood was collected from each individual and standard FicollPaque procedure was used to isolate peripheral blood mononuclear cells from freshly collected blood, which was diluted in a ratio of 1:1(v/v) with sterile phosphate buffered saline (PBS) and layered on equal volume of Histopaque density 1077 so that two layers are not intermixed followed by centrifugation at a speed of 300×g for 30min at room temperature in a swinging bucket without brakes. PBMCs ring was aspirated carefully through the tube with micropipette. PBMCs were further washed with PBS for minimum five times to ensure the platelets/plasma protein contamination free PBMCs pellet obtained by centrifugation at 200g for 10min.

### Extraction of proteins from isolated PBMCs

PBMCs were suspended in 300μL lysisbuffer(7M Urea, 2M Thiourea, 1% DTT, 2% CHAPS, 0.8% Pharmalyte (pH 3-10) and 10μg/mL Protease Inhibitor cocktail) and vortexed for 1min. followed by sonication for 15min. on ice. The lysate was centrifuged at constant speed of 12,000×g for 30min. at 4°C. Proteins in the supernatant were collected and stored at −80°C till further use for proteomics analysis by 2D-PAGE.Protein content in the supernatant was quantified with the help of Protein assay kit from Pierce Thermofisher as per the manufacturer’s instructions.

### Two dimensional polyacrylamide gel electrophoresis of PBMCs proteins

In order to nullify the inter-individual variations within the groups; equal amount of proteins (50μg from each) from at least five different subjects belonging to same group were pooled together and quantified and processed for 2D-PAGE.

In gel rehydration of IPG strip (7cm, pH3-10) was carried out with 60μg pooled protein in a final volume of 130μL rehydration buffer (8M Urea, 2%CHAPS, 0.002%Bromophenol Blue, 3mg/ml DTT and 1μL IPG carrier buffer (pH3-10) for overnight. Isoelectric focusing was carried out in Ettan IPGphore apparatus (GE Healthcare Life sciences, UK) for a total of 7500Vat a maximum of 50μA current and 20°C plate temperature over a period of 7.5h.Step1 100V for 4hours, Step2 step 300V for 1 hour, Step3 gradient 1000V for 30minutes, Step4 gradient 5000V for 1.5hours, Step5 step and hold 5000V till 2000Vh.After IEF, strips were first equilibrated in SDS equilibration buffer (6M Urea, 2%SDS, 0.375M Tris, pH 8.8, 20% glycerol) containing 130mM DTT for 15min and then DTT was replaced with 135mM iodoacetamide for next 15min under shaking conditions at room temperature.

### Sodium dodecyl sulphate polyacrylamide gel electrophoresis

Focused proteins in the IPG strip were then separated on 12.5% polyacrylamide gel. Proteomic profile of cell lysate was obtained by second dimension analysis on SDS-PAGE (11). Gels were stained with silver nitrate and analysed.

### Imaging and software analysis

Images were acquired by scanning gels in GE Ettan scanner and were saved in tiff format. Comparison of different groups was made with the help of Image Master Platinum 6 software (GE). Any artifacts if recognized as spots were manually checked and removed from the image data. Total number of spots, spot area and spot intensities were compared with the master gels of respective groups.

### Protein Identification by mass spectrometry

In gel digestion was carried out according to the following protocol. For silver stained gels, destaining of the differentially expressed spots was performed prior to tryptic digestion. Detailed protocol included spot picking from the 2D gel followed by cutting gel pieces (1–1.5mm) on a clean glass slide before destaining. Destaining was done with 1:1 mixture of 30mM Potassium ferricyanide and 100mM sodium thiosulphate followed by 4-5 washes with double distilled ultrapure water. The gel pieces were then dehydrated with ammonium bicarbonate in 40% acetonitrile for 30 min. at 37°C. Trypsin (0.4μg/10 μl) was used for in-gel digestion. Gel pieces were entirely covered with trypsin solution and incubated overnight at 37°C. Digested peptides were extracted from the gel pieces in the solution by brief sonication at 20% amplitude for 10 seconds, which were then used for MALDI-MS analysis.

### Mass spectrometry

Digested samples were desalted and concentrated on C-18 ZipTips (Millipore, Billerica, MA, USA) using the manufacturer’s protocol. ZipTips were eluted on MTP 384 target plate with 2μl of a-cyano-4-hydroxycinnamicacid (HCCA) (Sigma-Aldrich, USA) saturated solution dissolved in 50% ACN and 0.2% TFA. Mass spectra of digested proteins were acquired using Autoflex II TOF/TOF 50 (BrukerDaltonik GmbH, Leipzig, Germany) in positive reflectron mode, in the detection range of 500-3000m/z. External calibration to a spectrum, acquired for a mixture of peptides with mass range from 1046 to 2465Da, was done prior to acquisition. The proteolytic masses obtained were then processed through Flex Analysis v.2.4 programme for peak detection of proteins.

### Protein identification by Peptide mass finger printing

Peak detection in MALDI spectra and submission of peak lists to the Peptide mass fingerprint (PMF) to Mascot server were done using the Mascot Wizard program (Matrix Science, U.K). Peptide mass tolerance was set to 100 ppm with carbamidomethylcysteine set as fixed modification, oxidation of methionine as variable modification and 1 missed cleavage site was allowed. The peptides with high signal to noise ratio were preceded for MS/MS analysis and confirmed by matching with the *Homo sapiens* database.

### Bioinformatics and Protein-protein interaction/pathway analysis of identified proteins

The proteins that were identified by MS/MS analysis were further analyzed for their known functions, cellular location and their interaction with other proteins employing different bioinformatics tools. Functional characterization of the proteins was done using the SwissProt and Uniprot databases. The peptide sequences obtained from MS analysis were analysed for sequence similarity with the *Homo sapiens* and different organisms by using pBLAST program available at NCBI server (www.ncbi.nlm.nih.gov/BLAST/pBLAST). Physical and functional interactions between the differentially expressed proteins were predicted using the STRING (available at http://string-db.org/). Protein-protein interaction network analysis was performed by using Cytoscape 3.8.0. Properties of the network including node degree and edge attributes were then analyzed. Nodes represent proteins and edges represent the interactions/connections between the proteins. The degree represents the number of interactions associated with the protein. The Network Analyzer and MCODE app in Cytoscape 3.8.0 was used to compute the degree and between-ness centrality of the network.

## Results

### 2D-PAGE analysis

In order to find out signature proteins alterations in the peripheral blood mononuclear cells (PBMCs) in associated tuberculosis and diabetes patients, proteomics 2D-MALDI/MS approach was applied. The anthropological parameters of the patients and healthy controls are given in Table 2. The protein spots obtained by 2D-PAGE analysis were analyzed by ImageMaster Platinum6 software. Comparison was drawn between the groups and subgroups and mean protein expression spots were detected in 2DE gel profiles.

**Table 2:** Demographic and anthropometric characteristics of study samples of different groups.

| Groups |  | Age (years) | Gender (%) |  | BMI(kg/m<sup>2</sup>); Mean±SD |
| --- | --- | --- | --- | --- | --- |
|  |  |  | Male | Female |  |
| Group 1 | Naïve untreated tuberculosis only (n=20) | 28.2±10.3 | 60 | 40 | 18.1±3.8 |
| Group 2 Diabetes only | diabetes subgroup 1 (HbA1c 6.5-7.5)(n=20) | 46.0±9.3 | 70 | 30 | 26.8±7.2 |
|  | diabetes subgroup 2 (HbA1c 7.5-8.5)(n=20) | 48.0±6.7 | 60 | 40 | 28.6±4.8 |
|  | diabetes subgroup 3 (HbA1c >8.5)(n=20) | 52.6±4.1 | 70 | 30 | 30.7±5.8 |
| Group 3 | Associated Tuberculosis and diabetes(TBDM) (n=30) | 48.3±8.4 | 73 | 27 | 19.2±4.5 |
| Group 4 | Healthy controls (n=20) | 26± 4.0 | 50 | 50 | 21±2.1 |

TB; tuberculosis, DM; diabetes, BMI; Body Mass Index, SD; standard deviation, %; percent.

Mean of 310±18 protein expression spots were detected in healthy group as compared with a mean of 295±21protein expression spots in the naïve TB only group, a mean of 282±16 protein expression spots in the DM subgroup 1, 274 ±23 DM subgroup 2, 278±19 DM subgroup 3. In TBDM group 276±21 spots were identified. The spots match report of all the groups and subgroups along with number of spots upregulated or down regulated are represented in Table 3. When the gels from the different groups were compared, total 18 protein expression spots(approximately 8.3%) were identified with significant changes in spot intensity (expressed as % spot volume of at least >2 fold difference) (Fig 1). The selected spots were assigned molecular weight and pI with respect to molecular weight markers and pI markers respectively (Fig 2). The differentially expressed spots were further processed for MS analysis.

**Table 3:** Comparative computational analysis of 2DE-PAGE data of human PBMCs proteome among different groups using ImageMaster software.

| Groups | No. of spots | No. of Matches | Up-regulated | Down-regulated |
| --- | --- | --- | --- | --- |
| Healthy | 310± 18 | - | - | - |
| Tuberculosis | 295± 21 | 79 | 2 | 8 |
| Diabetes (HbA1c 6.5-7.5) | 282± 16 | 88 | 9 | 4 |
| Diabetes(HbA1c 7.5-8.5) | 274± 23 | 87 | 9 | 5 |
| **Diabetes(HbA1c>8.5)** | 278± 19 | 92 | 9 | 5 |
| **Tuberculosis and diabetes(TBDM)** | 276± 21 | 89 | 13 | 6 |
| **TB vs TBDM** |  | 73 | 10(TBDM) | 5(TBDM) |
| **DM vs TBDM** |  | 82 | 14(TBDM) | 7(TBDM) |

**Figure 1:**
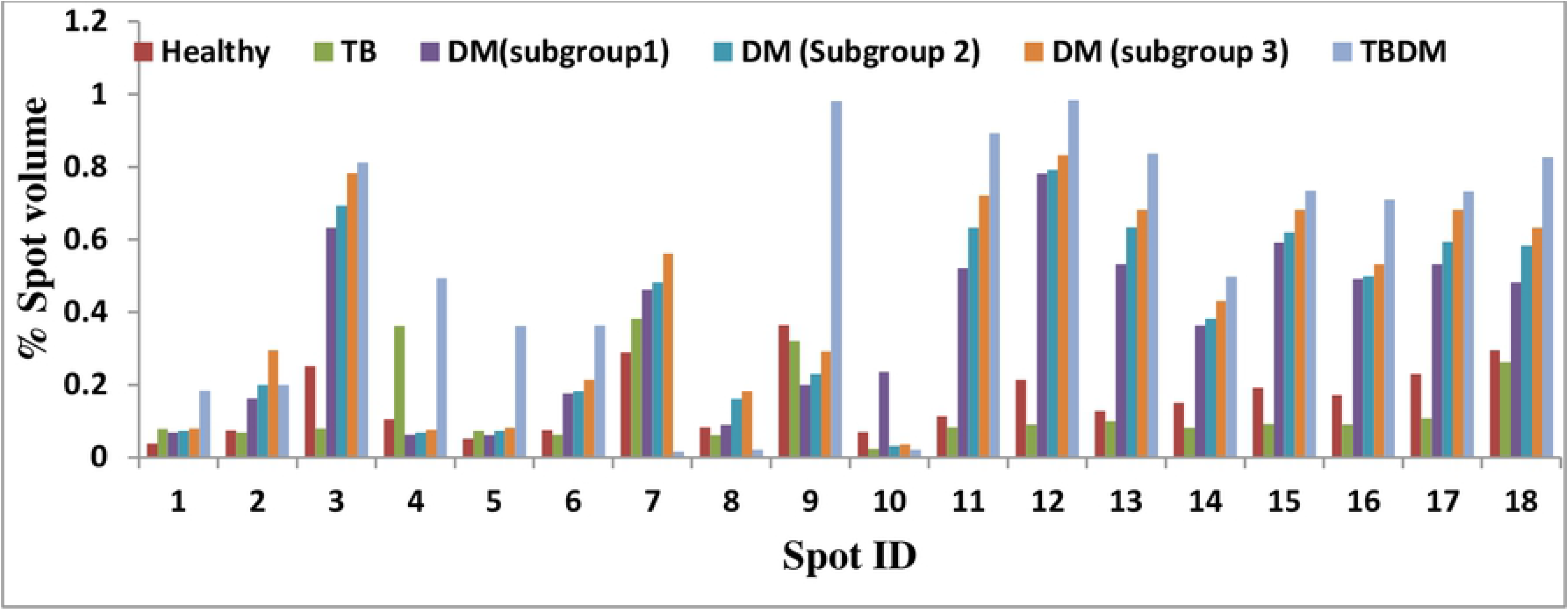
Histogram reflecting differential expression of PBMCs proteins in terms of spot intensity. (expressed as % spot volume of at least >2 fold difference; as compared to healthy controls.

**Figure 2:**
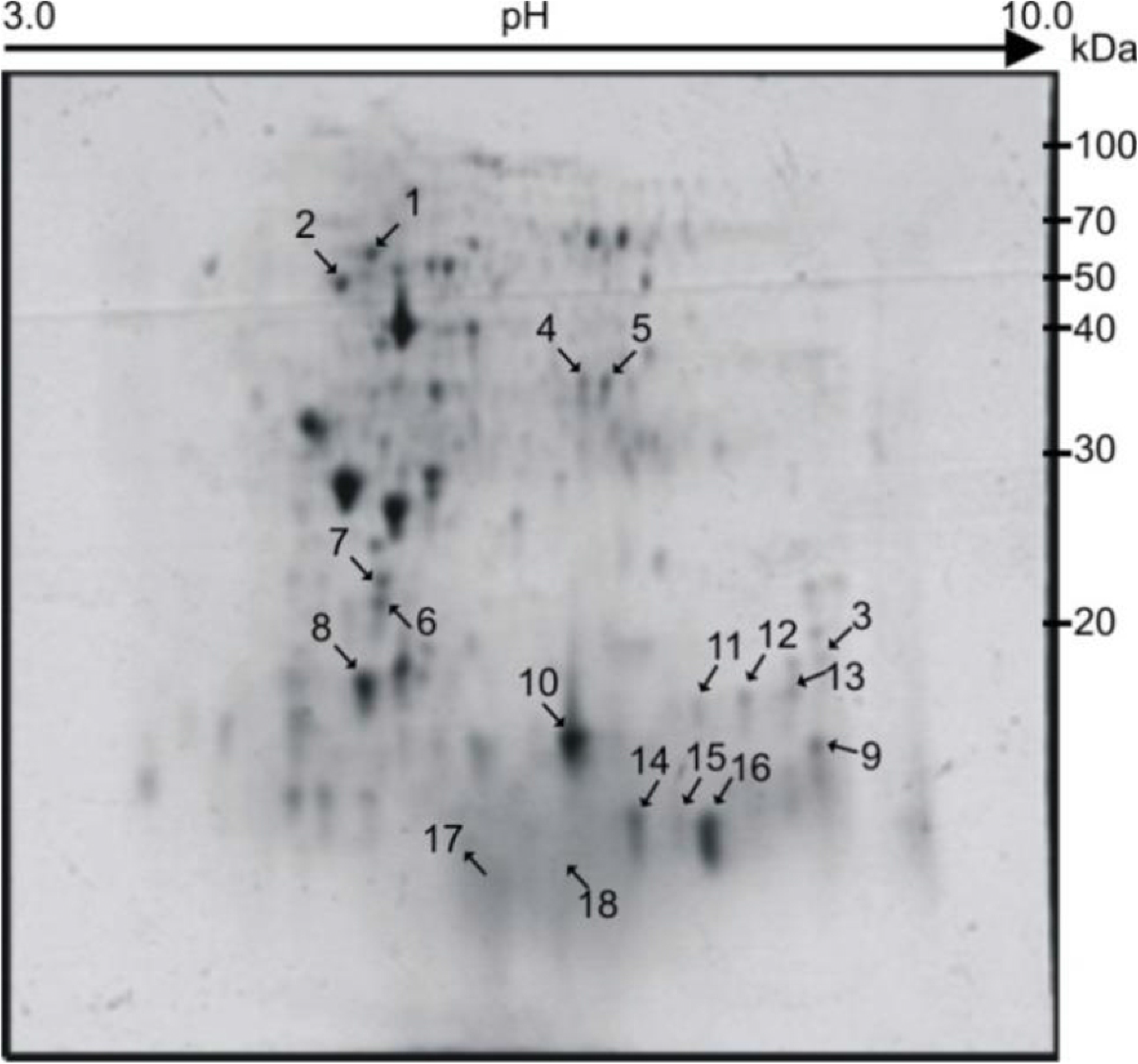
2DE-PAGE image showing PBMC protemic profile. Differentially expressed spots with corresponding spot ID are shown by arrows. At the top isoelectric point scale is shown (pH 3-10) from cathode to anode end of the IPG strip.

### Mass spectrometric analysis by Matrix Assisted Laser Desorption and Ionization-Time of Flight (MALDI-TOF)

Total 18 selected spots (found to be expressed in all groups and subgroups) were excised from the gel and overnight in gel trypsin digestion was carried out for each spot. The digested peptides were then spotted on MALDI target plate for MALDI-TOF mass spectrometric analysis. The mass spectrum along with m/z and intensity values was obtained for all spots followed by protein identification using MASCOT software. The identified proteins were analyzed for their subcellular location and function (Table 4) within the PBMCs using Swissprot, UNIPROT and NCBI databases. pBLAST analysis was performed for each identified protein. All the differentially expressed proteins showed 100% similarity with *Homo sapiens* (Table 5).

**Table 4:**
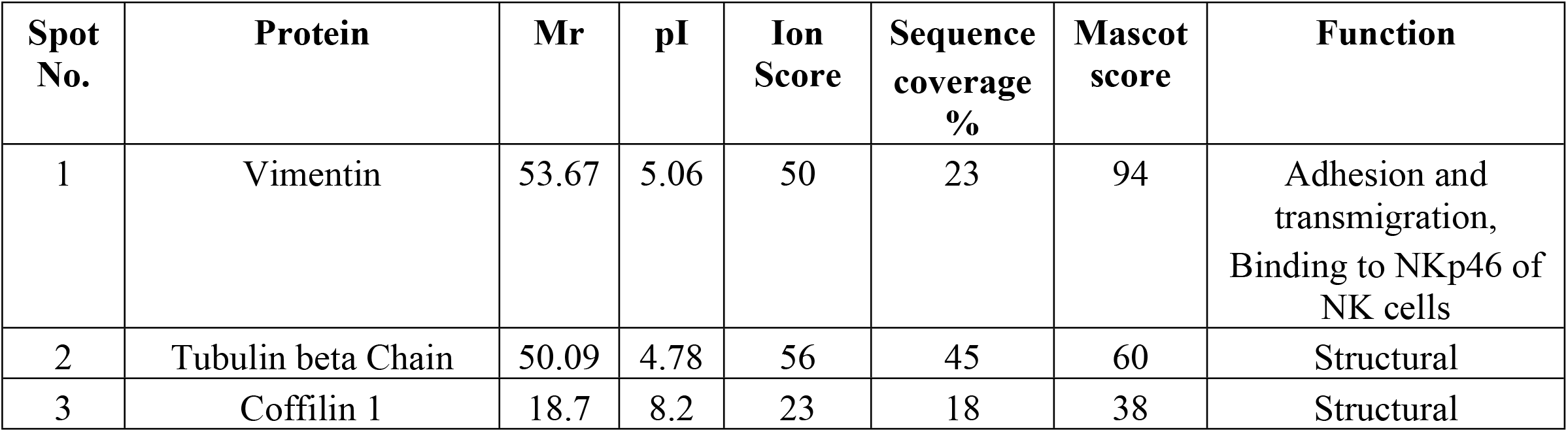
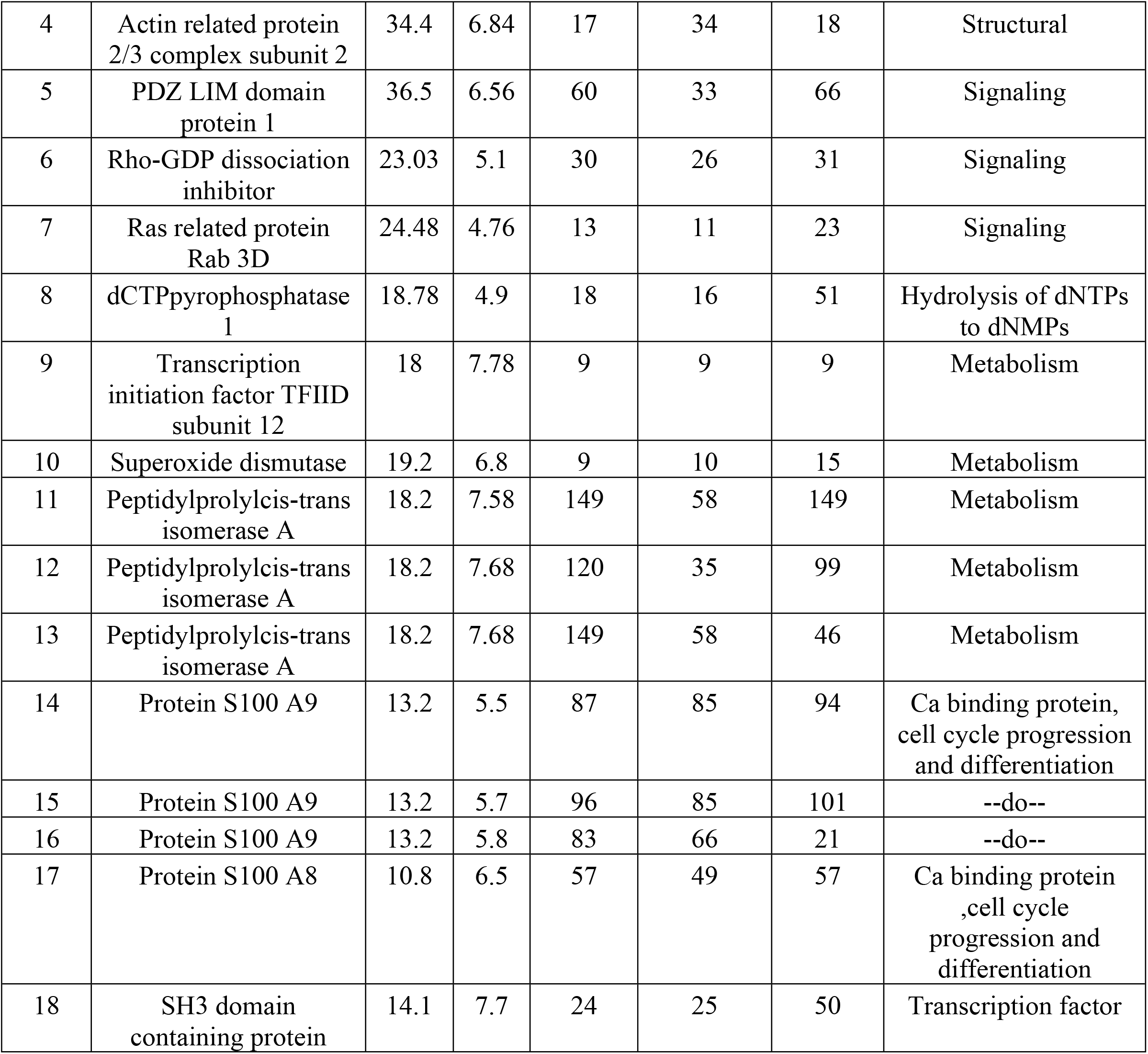
Detailed attributes and functions of the differentially expressed proteins identified by MALDI-TOF mass spectrometry.

**Table 5:**
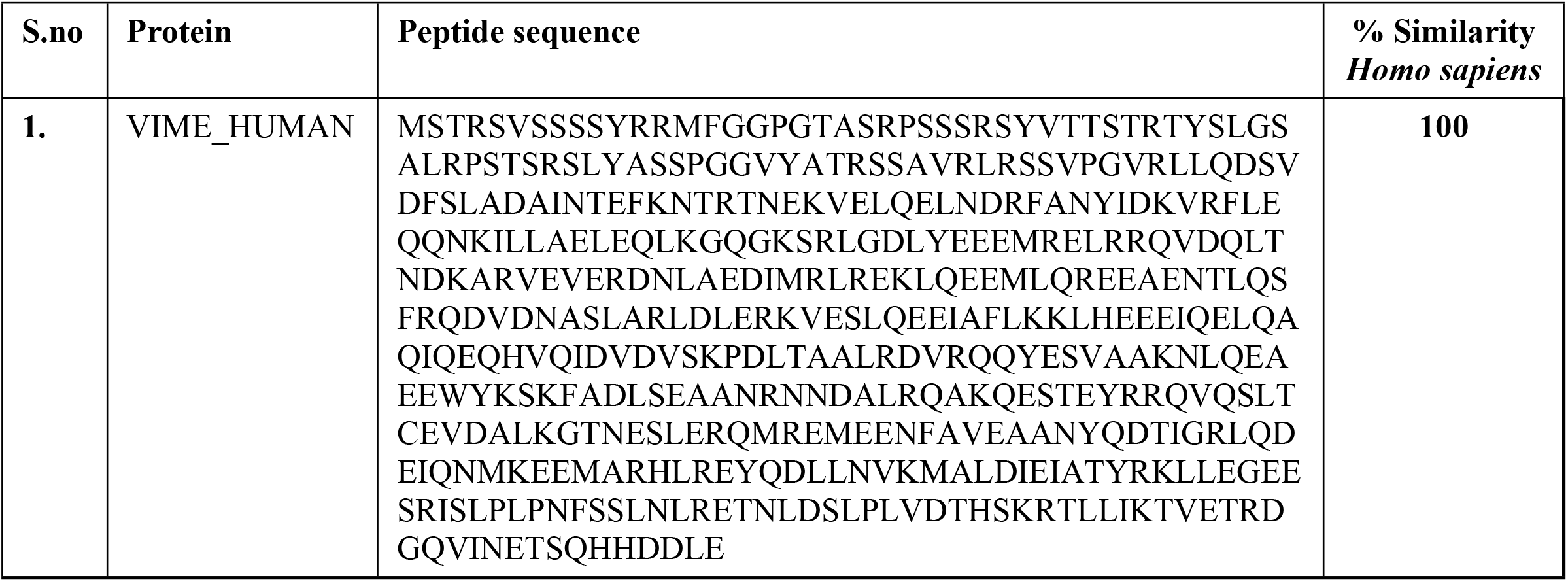
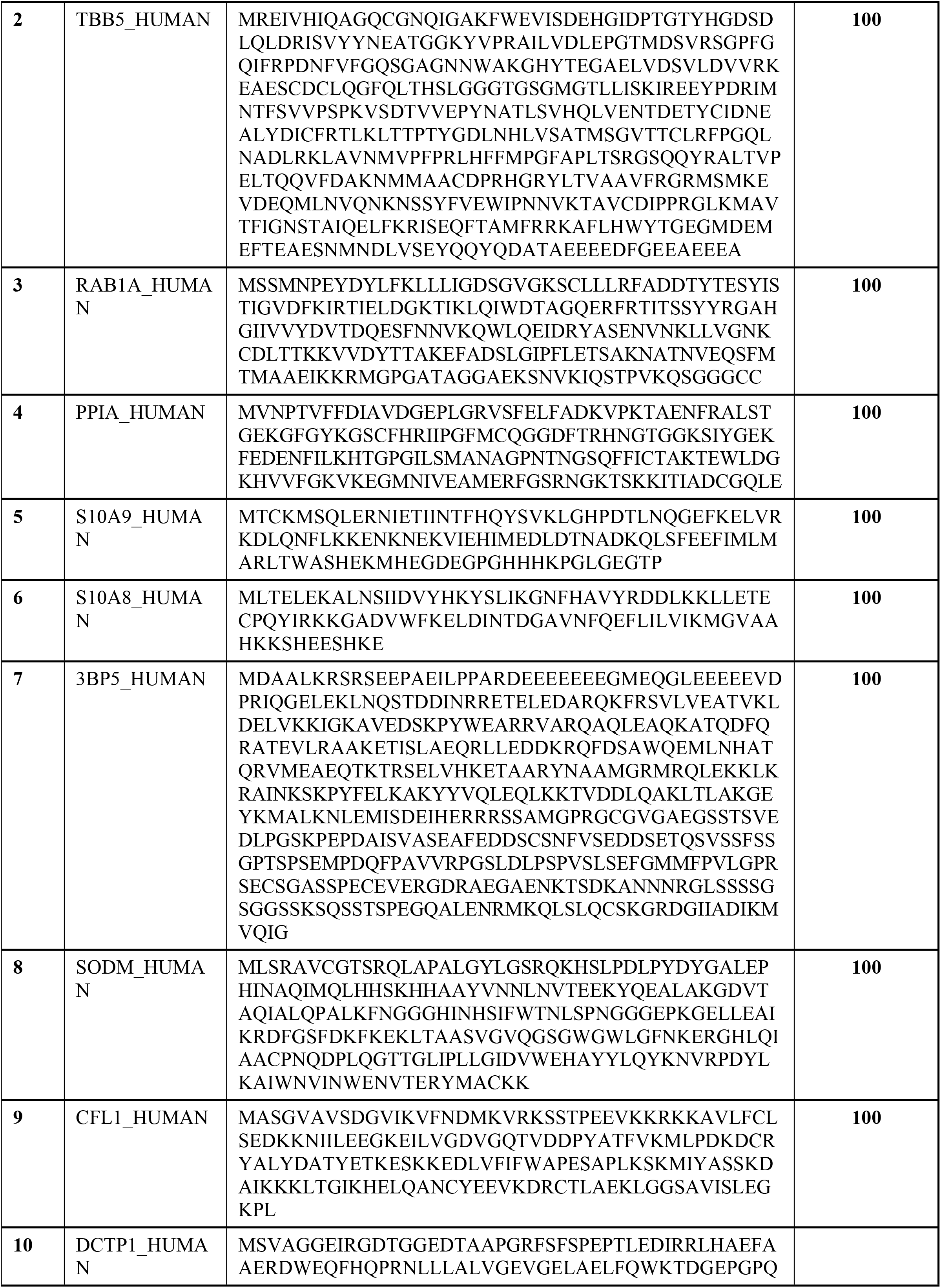
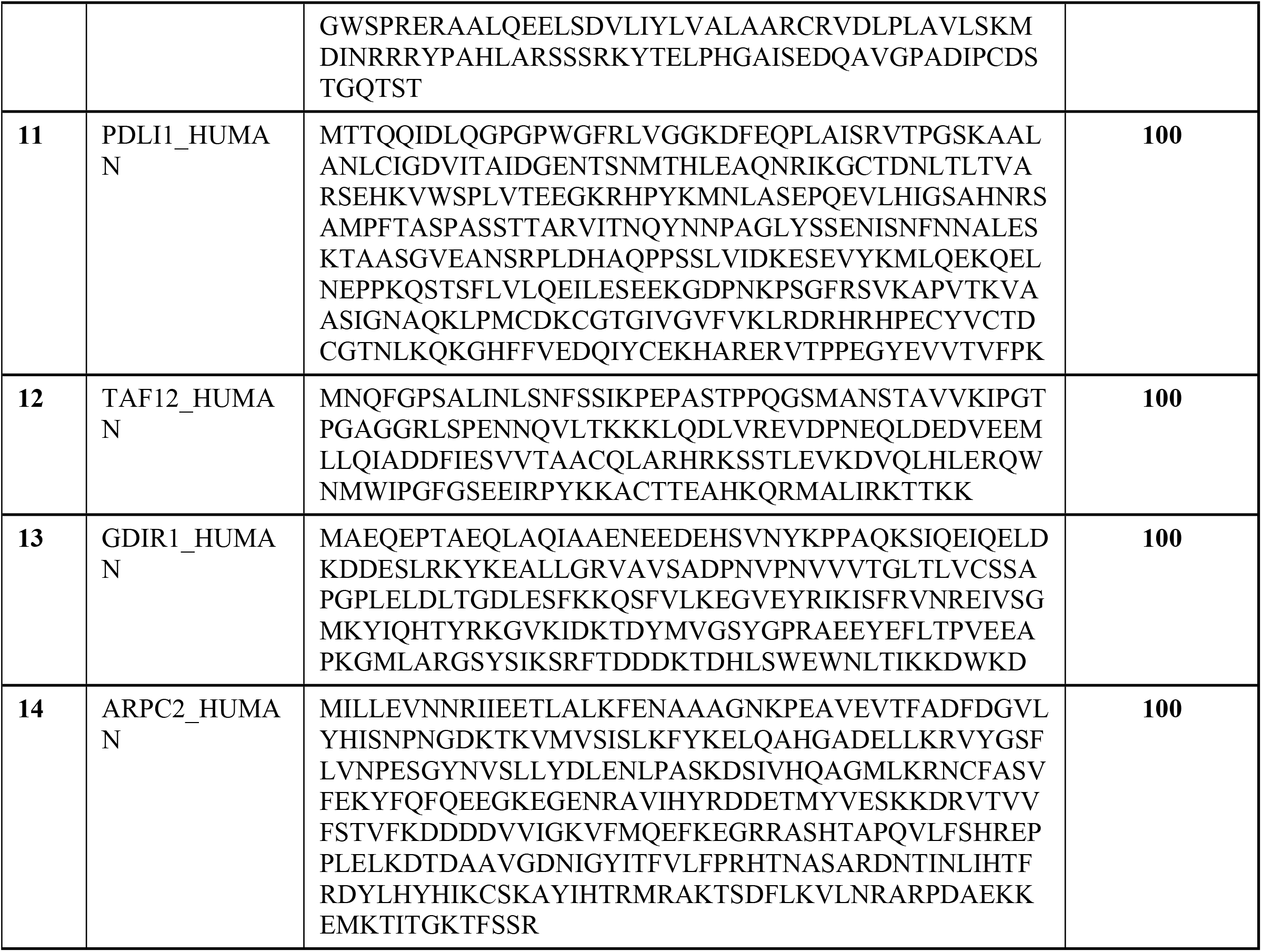
BLASTp analysis of identified peptide sequence from MS/MS analysis.

### Protein-protein interaction/pathway analysis

Protein-protein interaction analysis for the differentially expressed proteins was performed by using online softwares STRING and Cytoscape (Fig 3 and Table 6).

**Figure 3:**
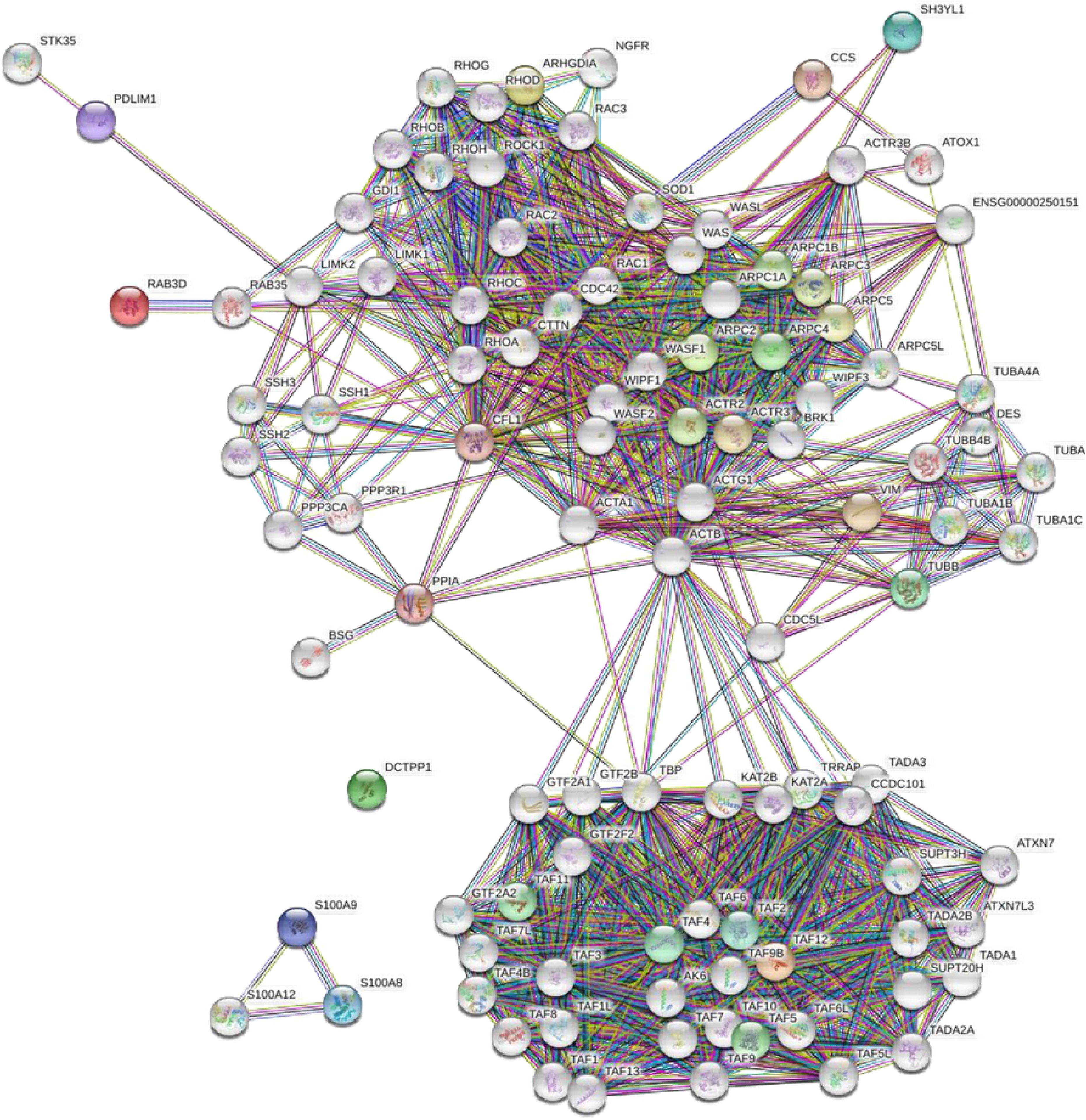
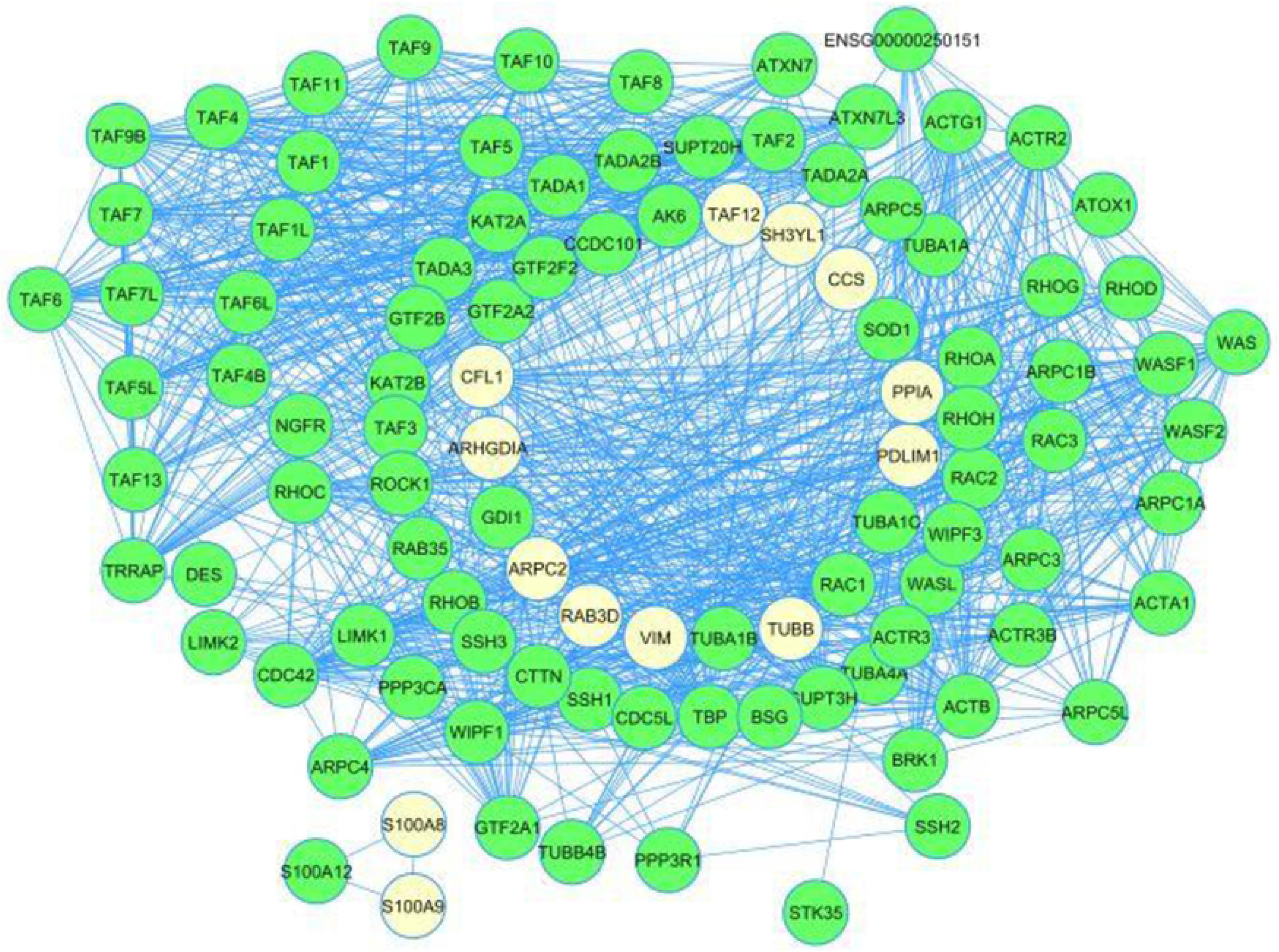
Protein-protein interaction network/pathway analysis by. **(a) STRING:** Different color nodes in the STRING image represents the individual proteins involved in different networks **and (b) module analysis by Cytoscape 3.8.0:** In the Cytoscape network proteins are represented as nodes, and the biological relationship between two nodes is represented as an edge (line). The most significant module in the protein-protein interaction network of differentially expressed proteins Module analysis of cytoscape software (degree cutoff=2, node score cutoff=0.2, K-core=2, and Max. depth= 100). White nodes in the cytoscape network represented differentially expressed proteins.

**Table 6:** Node data analysis using network analyser 4.4.5 from Cytoscape 3.8.0.

| S.No. | Protein name | Betweenness Centrality | Closeness Centrality | Clustering Coefficient | Degree | Neighborhood Connectivity | Number Of Directed Edges |
| --- | --- | --- | --- | --- | --- | --- | --- |
| 1 | ARPC2 | 0.004954 | 0.475962 | 0.809231 | 26 | 28.57692 | 36 |
| 2 | TAF12 | 0.001898 | 0.409091 | 0.793651 | 36 | 29.22222 | 2 |
| 3 | S100A9 | 0 | 1 | 1 | 2 | 2 | 2 |
| 4 | S100A8 | 0 | 1 | 1 | 2 | 2 | 26 |
| 5 | ARHGDIA | 0.006448 | 0.462617 | 0.783626 | 19 | 28.31579 | 19 |
| 6 | TUBB | 0.006535 | 0.5 | 0.681818 | 12 | 24 | 12 |
| 7 | CFL1 | 0.063619 | 0.529412 | 0.497561 | 41 | 24 | 41 |
| 8 | PPIA | 0.039479 | 0.492537 | 0.392857 | 8 | 23.875 | 8 |
| 9 | CCS | 0 | 0.314286 | 1 | 2 | 8.5 | 2 |
| 10 | VIM | 0.014356 | 0.428571 | 0.321429 | 8 | 20.75 | 8 |
| 11 | PDLIM1 | 0.020202 | 0.280453 | 0 | 2 | 7 | 2 |
| 12 | RAB3D | 0 | 0.298193 | 0 | 1 | 5 | 1 |
| 13 | SH3YL1 | 0 | 0.344948 | 1 | 3 | 36.66667 | 3 |

## Discussion

Tuberculosis and diabetes association is known since ancient times but its mechanism of association is not clear. A proteomics approach on the human PBMCs through 2DE-MALDI/MS tools was studied in order to find signature proteins related to copathogenesis of tuberculosis and diabetes. Unlike TB and Diabetes alone, body mass index of the copathogenic individuals was not associated with the disease severity. Further, in this study we obtained a modulated protein expression2DE proteomic profile in TBDM patient’s peripheral blood monocytes (PBMCs) in comparison to TB only, DM only pateints and healthy controls. Spot intensity comparison were made among healthy, TB only, Diabetes subgroup 1 (HbA1c 6.5-7.5), Diabetes subgroup 2 (HbA1c 7.5-8.5), Diabetes subgroup 3 (HbA1c >8.5) and TBDM groups.

A total of 18 mutually inclusive proteins were found to have differential expression and were present in all the groups as identified by MALDI-MS. According to their cellular functions annotated in Swiss-Prot, these differentially expressed proteins were classified into four categories i.e. structural proteins, signaling proteins, cellular metabolism intermediates and cell cycle and growth regulatory proteins. Many studies have shown that **Vimentin** plays an important role in adhesion and transmigration of infected monocytes and binding to NKp46 receptor of natural killer (NK) cells and these NK cells then lyse the infected macrophages (12, 13). In our study its expression was down regulated in TBDM patients as compared to DM only patients whereas in comparison to TB only cases its expression was found to be up regulated in TBDM cases. **Actin related protein 2/3 complex subunit 2** (Arp2/3) is a structural protein and helps in cell migration and phagocytosis activity. Phagocytosis is a hallmark of anti-bacterial host defense. Once the pathogens such as bacteria bind to PRRs, intracellular signaling pathways are triggered inducing actin polymerization. The process occurs through RacGTPases, and WASP (Wiskott-Aldrich syndrome protein) interaction which in turn activates Arp2/3 and actin polymerization occurs for phagosomal cup formation. *M. tuberculosis* impairs phagosomal maturation and this process is associated with actin nucleation followed by actin polymerization on endosomal membranes by activating the Arp2/3 complex. If this actin coat from the endosome carrying the mycobacteria is removed, mycobacteria will be delivered into phagolysosomes (14, 15). **Tubulin beta chain protein** is a main cytoskeleton protein that takes part in various cell movements during various cellular activities like migration and cell division. Its expression was enhanced in both DM only and combined TBDM cases in comparison to both TB only and healthy control group. **Rab1 small GTP-binding protein** plays a regulatory role in cell surface trafficking of human calcium sensing receptor (16). Another study on proteomic analysis of endocytic vesicles has shown that Rab1a regulates trafficking of early endocyticvesicles (17) and the level of Calcium sensing receptor (CaSR) present at the cell membrane. This protein also plays a role in cell adhesion and migration and *via* its role in protein trafficking also helps in autophagosome assembly and cellular defense reactions against pathogenic bacteria (16–18). Its expression was found to be down regulated in TBDM group in comparison to both only TB and only diabetes cases. **Peptidyl**-**prolylcis**-**trans isomerase** (PPIases) a cyclosporin A-binding protein cyclophilin (19) and its expression was upregulated in TBDM patients in comparison to only TB or diabetes patients. It is known to play a role in cyclosporine-A mediated immunosuppression in various pathologies. **Super-oxide dismutase (SOD)** was found to be upregulated in the DM and TBDM patients in comparison to TB only and healthy controls. SOD levels are found to be low in tuberculosis patients and higher in DM patients in comparison to healthy controls (20, 21). Upregulated expression of SOD has also been observed in *Helicobacter Pylori* infection, which supports the fact that the pathogen has developed protective mechanism against immune system (22). Similar protective mechanism might be adopted by *M. tuberculosis* in the copathogenesis of TBDM. **Protein S100A9** also called as calgranulin B was seen to be highly expressed in DM and TBDM group in comparison to TB only group. S100A9 have both antibacterial and antifungal activity (23). S100A9 always occurs in a heterodimer form with another protein S100A8. The dimer S100A8/A9 is commonly called as calprotectin and is constitutively expressed in myeloid cells. The dimer acts as a ligand of receptor for advanced glycated end products and toll like receptor 4. They are a part of endogenous inflammatory mediators called as Danger associated molecular patterns (DAMPs) (24, 25). In a study, it has been proved that S100A8/A9 proteins cause Neutrophil mediated inflammation in lungs during tuberculosis pathology (26). S100A9 has been recently predicted as an early diagnosis serum biomarker for pulmonary tuberculosis, inflammatory processes and immune response (27). It induces neutrophil chemotaxis and adhesion (28). The cumulative research supports our findings and it becomes pertinent to propose that levels of S100A9/S100A8 can be a potent biomarker for tuberculosis and diabetes copathogenesis. **PDZ LIM domain protein 1** expression was found to be enhanced in TBDM in comparison to both TB only and DM only patients. Some pathogen effector molecules and PDZ interactions are found to enhance bacterial spread in the mammalian cells through regulation of PKC pathway (29). PDLIM1 negatively regulates NF-κB-mediated signaling in the cytoplasm of Innate immune cells, such as macrophages and dendritic cell through sequestration of p65subunit of NF-κB thus reducing production of proinflammatory cytokine response to invading microbial pathogens(30).The role of the protein in connection with either of the two diseases i.e tuberculosis or diabetes is still not known.**SH3 domain-binding protein 5** is a negative regulator of BTK signaling in cytoplasm of B-cells and other lymphocytes. This protein was down regulated in the TBDM cases in comparison to the DM patients while its expression did not vary in comparison to only TB cases. **RHO protein GDP dissociation inhibitor of Rho proteins** (rho GDI) was found to be upregulated in diabetes group with HbA1c >8.5 and down regulated in TBDM patients in comparison to only TB patients in the present study. **TAFII subunit 12** is essential for mediating regulation of RNA polymerase transcription. Its expression was highly upregulated in the TBDM patients in comparison to both TB only and DM only patients.**Coffilin-1 protein** is actin-modulating protein. RHOGTPase negatively regulates its depolymerization activity by phosphorylating it through a downstream effector LIM kinase called as Rho-associated kinase (ROCK) (31). Cytoskeleton proteins and their regulation proteins could be influenced seriously in *M. tuberculosis* infection host cells leading to the apoptosis of host cells. These findings suggest that macrophages infected by *M.avium* could lead to apoptosis by regulating cytoskeleton protein β-actin or its regulatory protein cofilin-1 (32). Further bioinformatics analysis by STRING and cytoscape (Fig 3, Table 6) resulted in a common network in which all the differentially expressed proteins except DCTPP of the differentially regulated proteins were found to interact. These proteins make an interactome and work through a network for various cellular activites. BLASTp analysis of these peptides provided significant % sequence coverage of the protein and their sequences showed 100% similarity with the *Homo sapiens*.

## Conclusion

A set of 18 proteins were identified, which have modulated expression under copathogenesis conditions. The functional annotation to the identified proteins represents role of these proteins in the copathogenesis progression. Taken together, the identified proteins are found to be involved in modulating the host macrophages immunity and possibly provide mycobacterium tuberculosis a favorable environment to survive better. Three isoforms of two proteins named Peptidylprolylcis-trans isomerase A and Protein S100A9 were identified among the differentially expressed proteins which can be attributed to post translational modifications. These identified proteins may act as diagnostic or therapeutic targets to manage this comorbid situation in the host.

## Acknowledgments

We thank the patients for consenting to participate in the study.

